# Genome-wide regulatory model from MPRA data predicts functional regions, eQTLs, and GWAS hits

**DOI:** 10.1101/110171

**Authors:** Yue Li, Alvin Houze Shi, Ryan Tewhey, Pardis C. Sabeti, Jason Ernst, Manolis Kellis

**Affiliations:** Computer Science and Artificial Intelligence Lab, Massachusetts Institute of Technology 32 Vassar St, Cambridge, Massachusetts 02139, USA; The Broad Institute of Harvard and MIT, 415 Main Street, Cambridge, Massachusetts 02142, USA; Department of Organismic and Evolutionary Biology, Harvard University, Cambridge, Massachusetts 02138, USA; Department of Biological Chemistry, University of California, 615 Charles E Young Dr South, Los Angeles, California 90095, USA

## Abstract

Massively-parallel reporter assays (MPRA) enable unprecedented opportunities to test for regulatory activity of thousands of regulatory sequences. However, MPRA only assay a subset of the genome thus limiting their applicability for genome-wide functional annotations. To overcome this limitation, we have used existing MPRA datasets to train a machine learning model that uses DNA sequence information, regulatory motif annotations, evolutionary conservation, and epigenomic information to predict genomic regions that show enhancer activity when tested in MPRA assays. We used the resulting model to generate global predictions of regulatory activity at single-nucleotide resolution across 14 million common variants. We find that genetic variants with stronger predicted regulatory activity show significantly lower minor allele frequency, indicative of evolutionary selection within the human population. They also show higher overlap with eQTL annotations across multiple tissues relative to the background SNPs, indicating that their perturbations in vivo more frequently result in changes in gene expression. In addition, they are more frequently associated with trait-associated SNPs from genome-wide association studies (GWAS), enabling us to prioritize genetic variants that are more likely to be causal based on their predicted regulatory activity. Lastly, we use our model to compare MPRA inferences across cell types and platforms and to prioritize the assays most predictive of MPRA assay results, including cell-dependent DNase hypersensitivity sites and transcription factors known to be active in the tested cell types. Our results indicate that high-throughput testing of thousands of putative regions, coupled with regulatory predictions across millions of sites, presents a powerful strategy for systematic annotation of genomic regions and genetic variants.

## 1 Introduction

Disruption and aberrant coordination of gene expression is often at the upstream causes of complex human diseases. Genome wide association studies (GWAS) suggest the vast majority of disease-associated loci are located in non-coding regions, and are specifically enriched in enhancers, promoters, and other conserved cis-regulatory elements in disease-relevant cell types [1–8]. This implies that these regions harbor regulatory variants and are likely to be expression quantitative trait loci (eQTL). Systematically dissecting these GWAS risk loci requires characterizing the vast number of regulatory elements in the human genome, with a particular emphasis on those elements at which key transcriptional events occur that predispose to pathological changes. This in turn requires large-scale mapping of regulatory elements in multiple human tissues, an undertaking that has been the focus of large consortium efforts such as ENCODE [9] and the NIH Epigenomic Roadmap [10]. However, simply having a catalog of elements is not sufficient to understand how these elements work independently to regulate gene expression. This requires the ability to study elements in high throughput, and in the appropriate cellular context.

Until recently, large-scale experimental identification of candidate eQTLs and ultimately the disease-causing non-coding regulatory variants was infeasible. Computational approaches using supervised learning methods primarily focused on training models that predict more general disease causing variants from Human Gene Mutation Database [11], sequence specificities of protein binding sites [12], DNase I hypersensitive sites [13], and/or histone modification profiles [14]. However, even though each of these assays is individually informative, each of them only captures a partial view of regulatory function, they frequently disagree with each other, and they do not directly reflect the properties of cis-regulatory elements that drive gene expression.

To improve the ability to interpret non-coding sequence, approaches can directly measure the regulatory activity of a sequence in high-throughput are needed. One such solution is the massively parallel reporter assays (MPRA), which was recently developed to interrogate the regulatory potential of thousands of candidate sequences by measuring the down-stream activities of a reporter gene in a relevant cell type [15–22]. The assay is able to detect *regulatory hits*, which are the regulatory sequences that confer increased RNA expression relative to the basal promoter among thousands of ~150 bp candidate sequences. However, the assay has a relatively low sensitivity despite high specificity [21].

One approach is to examine the functional relevance of those features in explaining MPRA signals is to perform an explicit test on each individual feature to assess correlation with the regulatory hit. One approach is to examine the functional relevance of those features in explaining MPRA signals, and then perform an explicit test on each individual feature to assess correlation with the regulatory hits [18, 20, 23, 24]. Another approach is to train a classification model on predicting MPRA response using composite features. Even though MPRA assays only cover a small fraction of the genome, they present the opportunity to directly harness the resulting information in order to improve the ability to identify eQTL and GWAS causal alleles and to ultimately predict regulatory variants associated with complex human diseases. Moreover, comparison of the prediction accuracies within and between different MPRA datasets generated from varied experimental approaches may reveal further insights as to the effectiveness and differences of regulatory elements in predicting regulatory potential.

In this study, we formulate a supervised learning strategy to facilitate MPRA data analysis and then use those data to infer potential regulatory SNPs among all common variants. We demonstrate that the regulatory hits from MPRA data are highly predictable when both sequence and functional genomic data are taken into account. We identify a set of meaningful regulatory binding sites, motifs, and epigenomic features that are highly predictive of the MPRA outcomes (i.e., whether a sequence confers active *in-vitro* regulation of transcription or not) in a given cell type, and also indicative of the transfected cell line’s tissue of origin as shown by the DNase I hypersensitive sites assayed in the same cell type. Finally, we use this information to build a general model that scores 14.3 million common non-coding variants by predicted MPRA potential (PMP) in three distinct cell types.

## 2 Results

### 2.1 MPRA regulatory sequence predictions overview

In this paper, we used two recent MPRA datasets to train our model (Fig. 1a). For each dataset, we processed them into a set of positive examples and a set of negative examples.

**Figure 1:**
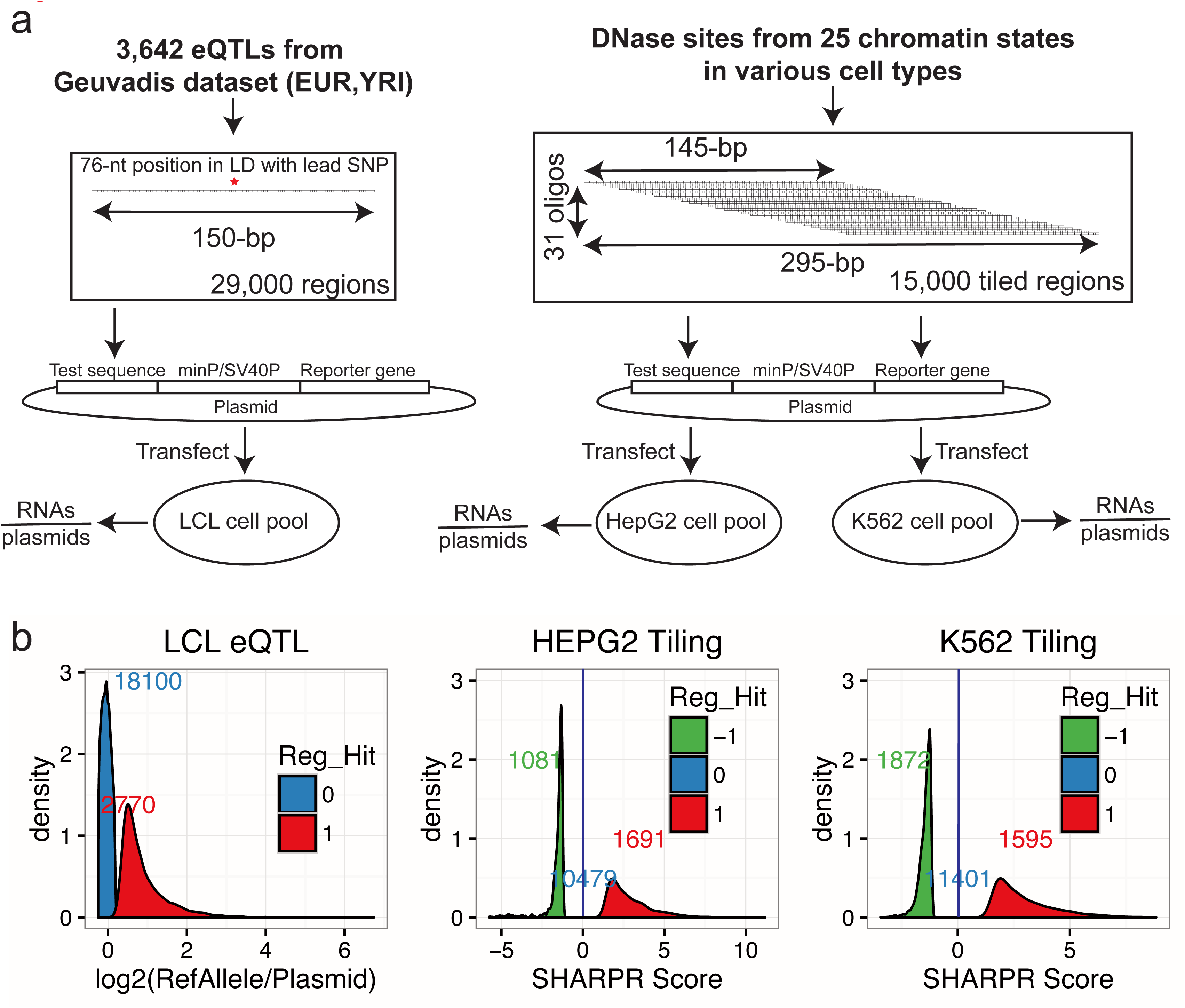
Overview of the MPRA computational analysis. (a) Experimental design and generation of the MPRA datasets used in this study: the LCL dataset derived from eQTLs (left) and tested in a lymphoblastoid cell line (LCL) and the the DNase-tiling dataset derived from 25 chromatin states and tested in HepG2 and K562 cells (right). After transfection of plasmids containing each regulatory sequence into appropriate cells, transcribed RNA was quantitated, and the fold-change ratio of RNA over plasmid was calculated to quantitate the extent of the corresponding sequence regulating transcription. For the tiling MPRA datasets, a sequence could be either activating or repressive whereas the eQTL-MPRA dataset only tests for activating regulatory hits in LCL. (b) Constructing confidence training data. We created confidence positive and negative training cases. The density plots display the corresponding distribution of each MPRA dataset and the total numbers of positive/negative training cases that went into the supervised training procedures. The x-axis for the two tiling datasets is the Sharpr-MPRA regulatory activity scores (SHARPR) computed from tiled reporter data [22].

The first dataset were generated from lymphoblastoid cell lines (LCLs), looking to validate eQTLs from the Geuvadis RNA-seq dataset of LCL from individuals of European and West African ancestries [21]. The candidate regulatory sequences were generated using 150-bp oligos centered at the lead SNP (or SNPs in perfect LD to the lead SNP) in 3,642 high-confidence eQTLs, and tested in LCL. To enable binary classification, we obtained only the activating sequences in the LCL-eQTL MPRA data that exhibit significantly higher fold-change of RNA over plasmid with Bonferroni-adjusted p-value < 0.01 and negative sequences with fold-change within one standard deviation from the average fold-change. As a result, we obtained 2,770 positive (i.e., activating regulatory hits) and 18,100 negative training cases (Fig. 1b).

The second dataset were derived from DNase-sensitive sites selected based on 25 ChromHMM chromatin states in four cell types [22]. The tiling MPRA datasets were generated by tiling 31 overlapping 145-bp sequences across 15,720 295-bp regions and tested in HepG2 and K562 cell lines. To generate training examples, we applied Gaussian mixture model on the inferred MPRA regulatory activities at each single-nucleotide by the previously published MPRA-SHARPR model. We then selected activating and repressive regulatory sequences based on posterior probabilities above 0.9 and 0.95, respectively, and negative control sequences within 1% standard deviation from zero. As a result, we obtained 1,691 and 1,595 activating, 1,081 and 1,872 repressive, and 10,479 and 11,401 negative controls for the tiling MPRA data generated from HepG2 and K562 cell lines, respectively (Fig. 1b).

For ease of reference, we will refer to the above MPRA training datasets as “LCL_act”, “HEPG2_act”, “K562_act”, “HEPG2_rep”, and “K562_rep”, respectively.

### 2.2 MPRA regulatory hits are highly dependent on cell-type specific DNase-sensitive sites

As features in our machine learning predictive model, we used reference genomic and epigenomic annotations. We compiled one of the largest set of functional features consisting of 3171 functional genomic features uniformly processed based on the GRCh37/hg19 genomic coordinates for each of those 150-bp training cases. These include:

- We used epigenomic annotations from 1,032 epigenomic signal tracks (H3K4me1, H3K4me3, H3K36me3, H3K27me3, H3K9me3, H3K27ac, H3K9ac, DNase I hypersensitive) estimated as −log p–values by MACS2 [30] across 127 cell types from the ENCODE/Roadmap Consortium [10];
- We also used ENCODE ChIP-seq transcription factor binding peaks for 161 transcription factors in various cell types (primarily from GM12878 and K562) as binary features from the UCSC database [9];
- In addition, we used sequence features corresponding to regulatory motif occurrences. We compiled datasets of 1,934 position weight matrices (PWM) for 602 transcription factors [25] and we searched their occurrences across the genome. This resulted in 1,934 occurrences, which we treated as binary feature for motif presence or absence for every one of these regulators;
- Moreover, because repeats have been widely hypothesized to be predictive of regulatory binding and often shown to be correlated with expression activities [ref]. We therefore also included 43 binary features for each of the repeat families from Repeat Maskers database from the UCSC database;
- 1,032 epigenomic signal tracks (H3K4me1, H3K4me3, H3K36me3, H3K27me3, H3K9me3, H3K27ac, H3K9ac, DNase I hypersensitive sites) estimated as −log p-values by MACS2 across 127 cell types from the ENCODE/Roadmap Consortium [10];
- Lastly, since evolutionary conservation is highly correlated with regulatory function, we also included evolutionary conservation track from phast46way placental conservation score obtained from the UCSC database.

We then sought to gain insights as to what the features that are most informative in MPRA predictions. We first investigated how well each individual feature correlate with MPRA regulatory hits to gain some biological insights on what dictate the MPRA outputs. The top scoring features in each MPRA training set were biologically meaningful in terms of the cell types and regulatory factors (Fig. 2). For example, DNase-sensitive sites were among the most significant features for each MPRA dataset (highlighted in Fig. 2). In addition, repressive regulatory hits in the tiling MPRA dataset were strongly enriched for transcriptional repressor REST and RFX5 binding motifs in HepG2 cells, as independently highlighted in the original paper [22].

**Figure 2:**
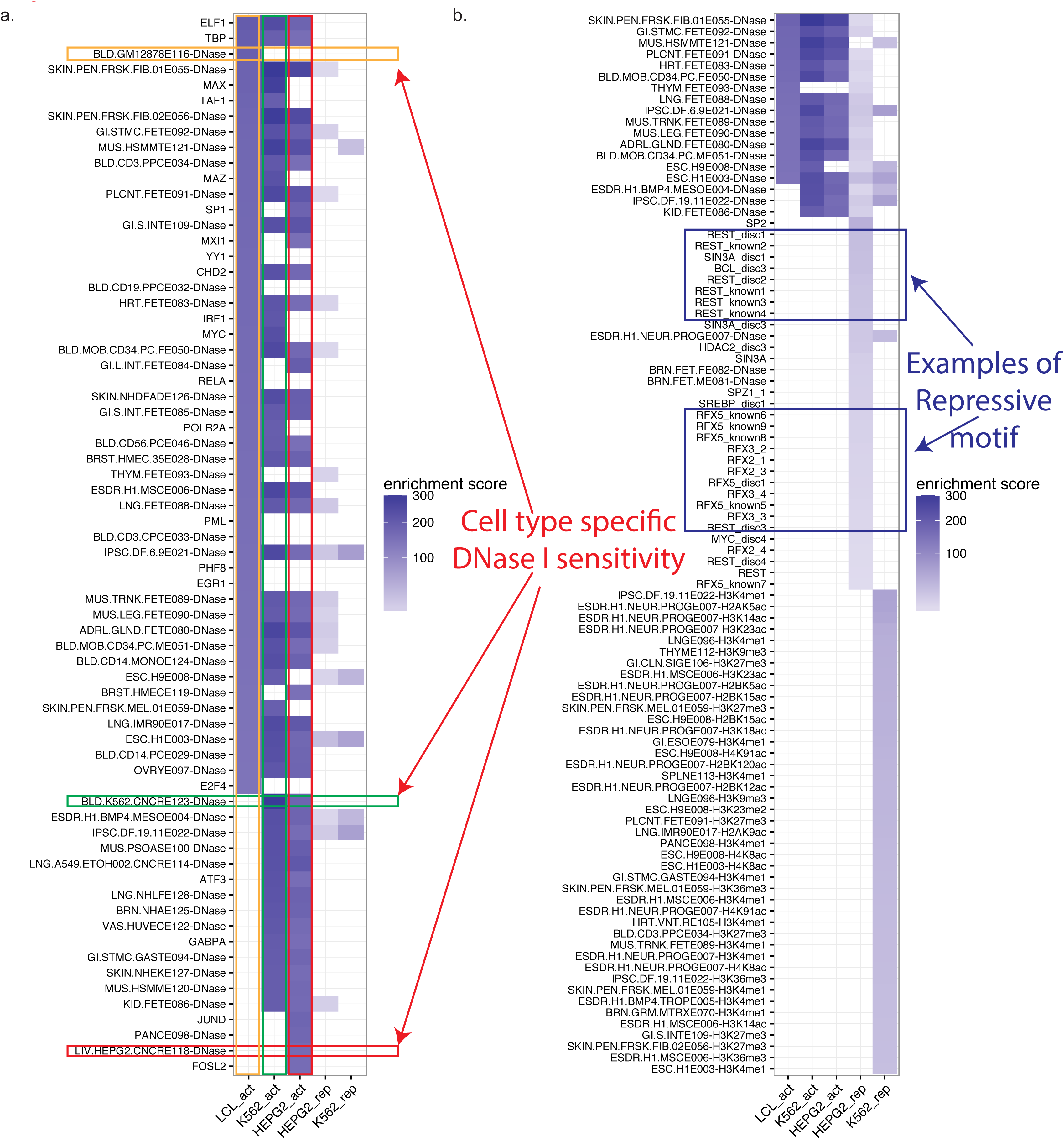
Feature enrichment. Top 50 features with the highest enrichment score for each of the five MPRA datasets. These features include epigenomic signals, ChIP-seq transcription factor (TF) binding, TF sequence motifs (Supplementary Table S1). The top features are divided into the features significantly enriched in the (a) transcriptionally activating MPRA hits (i.e., HepG2 act, K562 act, LCL act) and (b) transcriptionally repressive MPRA hits (i.e., HepG2 rep and K562 rep).

### 2.3 Sequence motifs are the predominant features in MPRA predictions

To further examine the underlying importance of each functional genomic feature, we prioritized the features based on the magnitude of the linear coefficients from the elasticnet model trained on all 3171 functional genomic features. Although the presence of a DNase-sensitive site was among the most significantly enriched features in the above single-feature test, known sequence motifs, repeat families, and TF binding sites exhibit the higher importance in the multiple regression elasticnet model (Fig. 3). Motifs are known to play important roles in expression regulation [18, 22, 25]. We further ascertained our results by comparing our motif rankings with published motif hits in the tiling array analyses (i.e., Supplementary Table 2 from [22]). We observed a significantly higher cumulative overlap between motifs ranked by the observed coefficients and the published MPRA motif hits [22] compared to a background constructed by permuting coefficients among the motifs (Fig. 4a). Most top motifs that we found are also active in both HepG2 and K562 cell lines (Fig. 4b).

**Figure 3:**
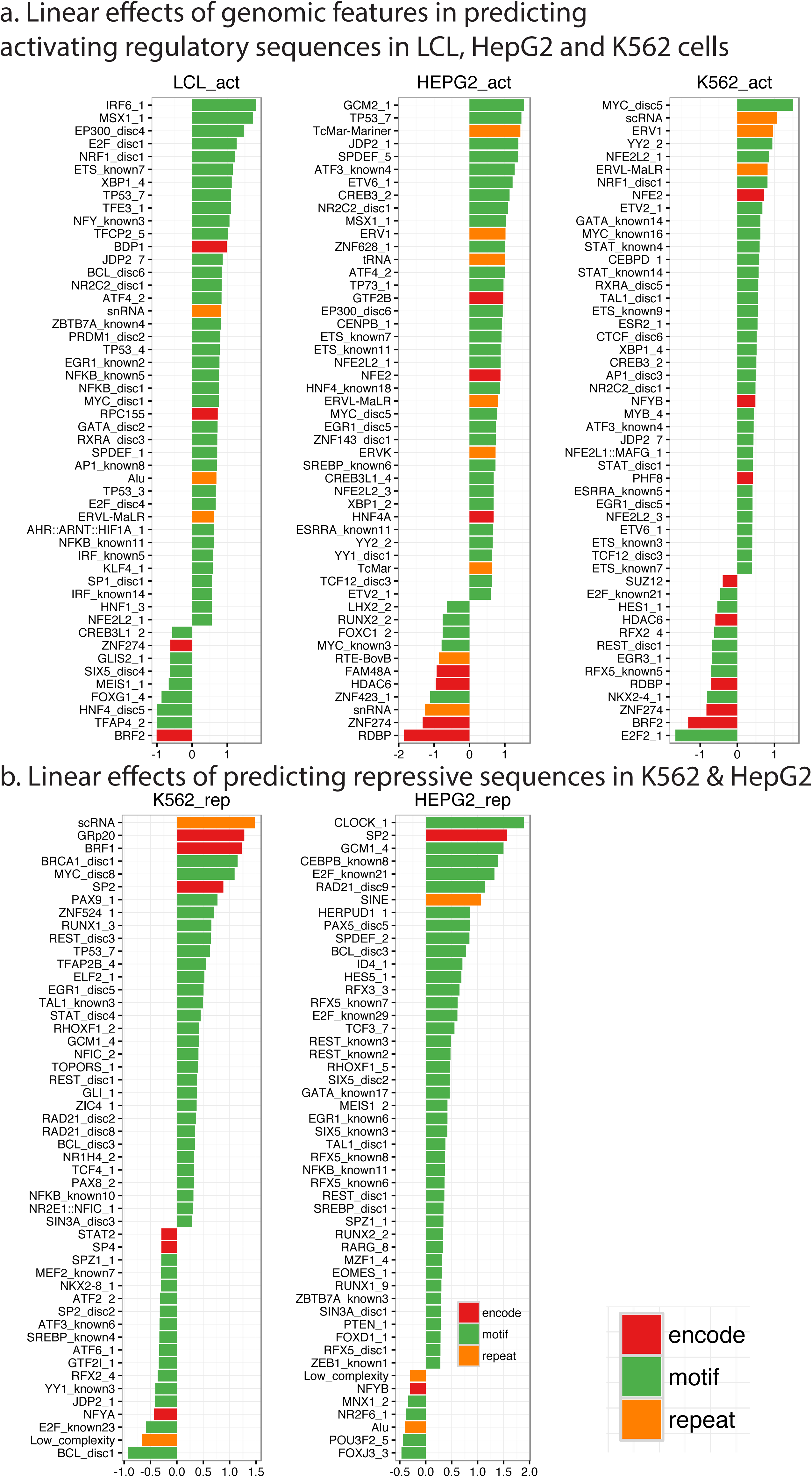
Top features in MPRA predictions. For each cell type and each experiment, features were ranked based on the corresponding linear coefficients that were fit by the elasticnet model. Colors indicate different functional categories including ENCODE TF bindings (encode), known sequence motifs (motif), and repeat families (repeat).

**Figure 4:**
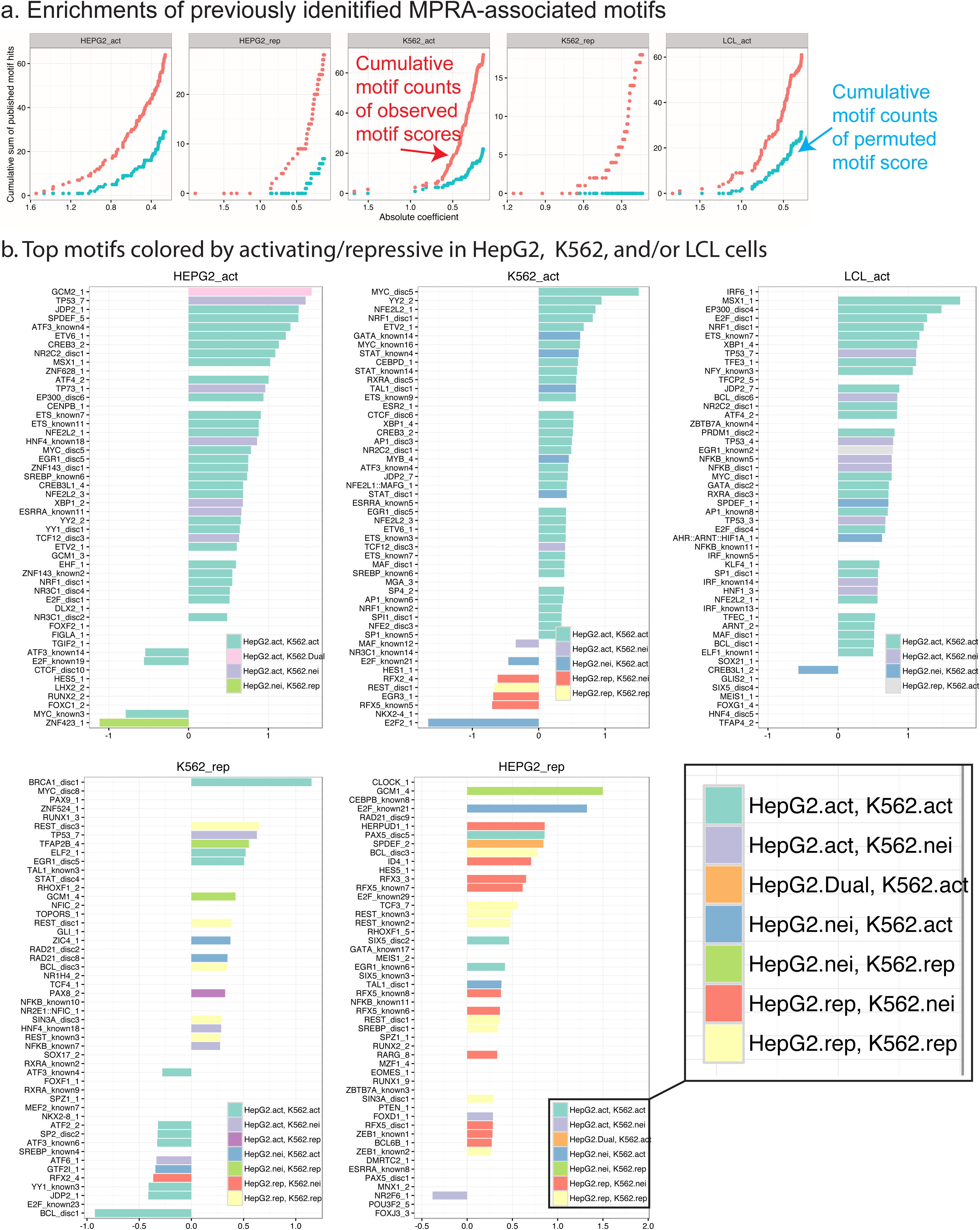
Comparison with known motifs from previous tiling MPRA analysis [22]. (a) Cumulative overlapped counts of published MPRA motif hits as a function of decreasing linear coefficients from the elasticnet model. Observed and permuted are colored in red and cyan, respectively. (b) Top hits that were also found in the known motif hits from previous study [22].

### 2.4 MPRA regulatory hits are highly predictable using functional genomic and sequence features

Importantly, we found that MPRA regulatory sequences are highly predictable in many cases based on functional genomic and sequence features, indicating that we can indeed make genome-wide predictions from those trained models. Specifically, we evaluated elastic-net using the 3,171 genomic annotations as features, gapped k-mer SVM (gkm-SVM), and elasticnet_gkm (i.e., using the 3,171 genomic annotations plus predictions from gkm-SVM as an additional feature) on their abilities to classify withheld data in a 22-fold cross-validation procedure, where each fold corresponds to a distinct chromosome. Most methods demonstrated comparable performances (Fig. 5). Elasticnet, the regularized linear model (i.e., regularized linear regression with L1/L2 norm), surpassed the sequence-based gkm-SVM in predicting activating regulatory hits from the tiling data. In particular, elasticnet achieved 90% AUC of ROC (AUROC) and 67% AUC of precision-recall (AUPRC) on the HepG2 tiling data and 93% AUROC and 72% AUPRC on the K562 data, compared to 88% AU-ROC and 63% AUPRC on the HepG2 tiling data and 91% and 67% on the K562 data from gkmSVM (Supplementary Table S2). Elasticnet-gkm did not improve performance over elasticnet in predicting tiling MPRA signals. On the other hand, elasticnet-gkm and gkm-SVM perform better than elasticnet on the non-tiling eQTL-LCL data. Thus, there is no single method that performs the best in all three datasets.

**Figure 5:**
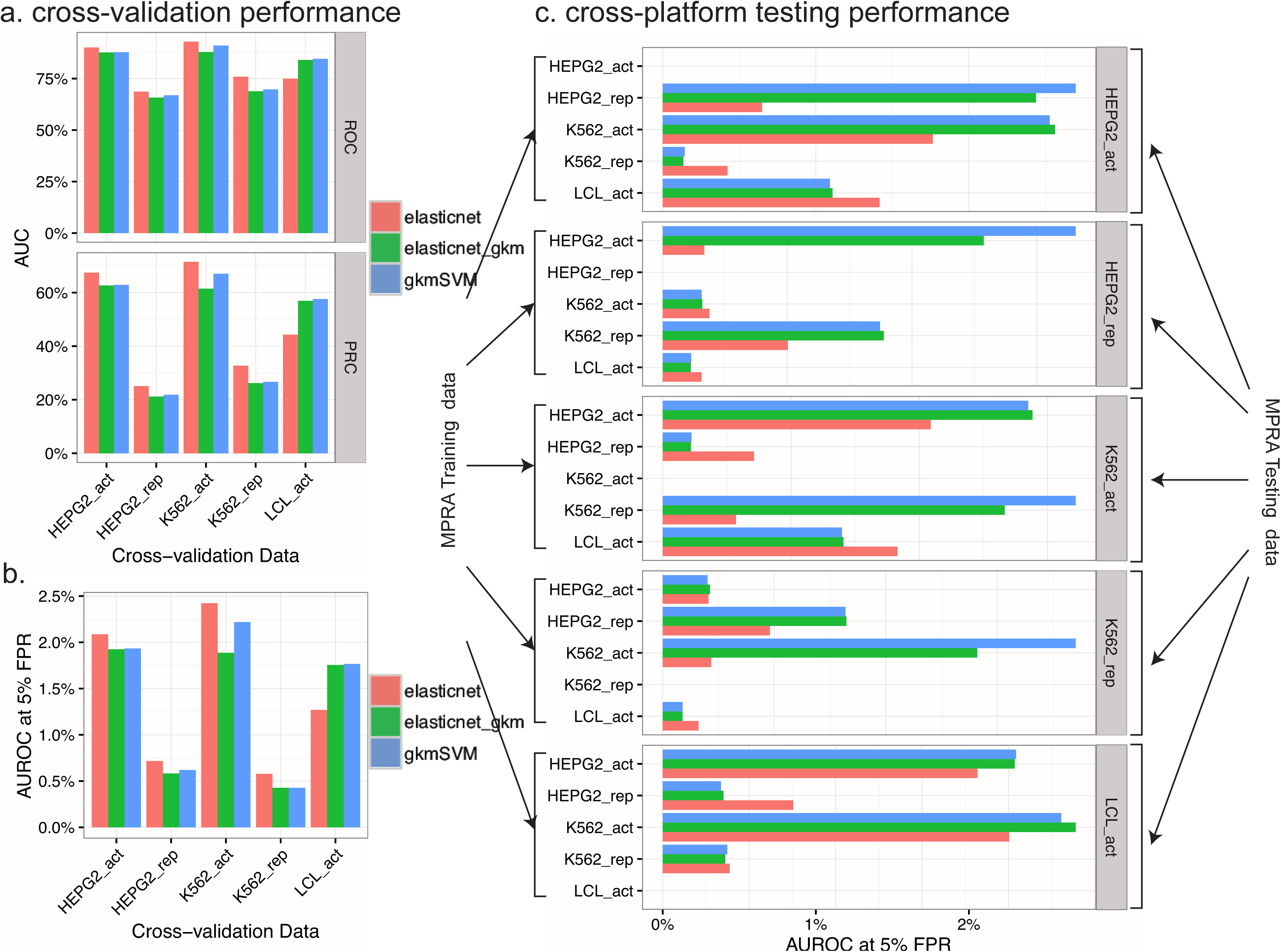
Power analysis. (a) Area under the ROC and PRC curves of each model for predicting regulatory hits in each of the five MPRA datasets. All performances were evaluated in cross-validation (Materials and methods). (b) AUROC up to 5% false prediction rate. Sensitivity is defined as the percentage of true positives out of all positive cases. (c) Testing performance. Each model trained on one MPRA dataset was tested on another MPRA dataset. The performance was assessed by AUROC up to 5% FPR to facilitate cross-platform comparison.

Due to the imbalance of positive and negative data within each dataset, AUPRC is provide additional important metric to evaluate the method performance. Also, the various MPRA training datasets contained different proportions of positive and negative training examples, which made cross-platform comparison difficult based on AUPRC. In this regard, we included AUROC up to 5% false positive rate (FPR) (Fig. 5b) and noted that the relative performance was qualitatively consistent with all metrics. On the other hand, repressive regions were much more difficult to predict, with lower performance among all methods possibly due to various reasons (Fig. 5a, b, **Discussion**).

### 2.5 Testing performance is platform and cell-type dependent

We found that our approach also works in cross-platform predictions, indicating that the trained model can generalize to unseen data. Specifically, we trained the three classification methods on one dataset and applied them to predict regulatory hits in another dataset. We compared the power of each trained model (method + training dataset) based on an AUROC up to 5% FPR (Fig. 5c) and summarized the results as follows.

1. The trained models conferred reasonable power when predicting unseen data in the relevant context. For example, elasticnet trained on HepG2_act data achieved close to 2% AUROC up to 5% FPR on predicting K562_act, which is slightly worse than the AUROC from the cross-validation of the same model on K562_act;
2. The elasticnet model trained on the 3,171 functional genomic data performed generally better on predicting unseen activating testing cases than on predicting repressive hits;
3. The gkm-SVM and elasticnet-gkm, using sequence features, conferred reasonable performance not only in predicting the same regulatory activity (e.g., trained and tested on activating data) but also on predicting opposite regulatory activity in the same cell type (e.g., trained on activating hits in HepG2 and tested on repressive hits in HepG2);
4. The performance is platform-dependent – models trained on tiling data (i.e., K562 and HepG2) performed better on predicting on tiling signals than they did on non-tiling signals (i.e., LCL);
5. Models trained on the blood cell-line K562 data performed slightly better than models trained on the liver cell line HepG2 data in predicting targets in the other blood cell-line LCL data and vice versa.

### 2.6 Scoring common variants using the trained MPRA models

We extended the utility of MPRA to infer genome-wide regulatory potentials, we trained our machine learning model on each of the five MPRA datasets and then use them to score all common variants. We observed discriminative power of our model in distinguishing common variants in the tested regulatory regions and a large number of untested common variants that exhibit promising regulatory scores. Specifically, We chose elasticnet-gkm as the representative model due to its competitive performance observed above. We then applied our model trained on the full MPRA datasets to score 14.3 million common variants from SNPdb (version 146) obtained from the UCSC browser, effectively constructing a novel genomic tracks indicative of regulatory potentials learned from the MPRA data (**Supplementary Table S5; Materials and methods**). We first verified that variants located within the training MRPA regulatory sequences exhibited significantly higher prediction scores than those of the remaining variants from each model (Fig. S3).

Moreover, we assessed the fraction of putative regulatory common variants with scores above the median prediction scores of the observed regulatory hits from each MPRA experiment. Overall, 0.21% and 2.6% of the common variants are predicted to be regulatory depending on the trained models and yet only lower than 0.0083% and 0.033% of the common variants are covered by any the current MPRA training regulatory sequences from either tiling and non-tiling MPRA data, respectively (**Supplementary Table S4**). Thus, potentially a large number of regulatory variants were untested by the assays.

### 2.7 Functional implication of predicted regulatory scores of common variants

Remarkably, the predicted regulatory scores are negatively correlated with the minor allele frequency of SNPs and are highly discriminative of SNPs harbored in eQTL from GTEx in various tissues, and SNPs reported in GWAS catalogue relative to background SNPs (Fig. S4, S5, S6). Specifically, we assessed the functional implications of the predicted MPRA potential (PMP) scoring system based on minor allele frequency (MAF), GTEx eQTL data (version 6) [26], and SNPs from GWAS catalog (v1.0 2016-01-22) [27]. Notably, none of these categories was present in the training data. We observed that the variants among the top 1% quantile of the predicted scores exhibited significantly lower MAF (FDR<10^−70^) than the remaining SNPs (Fig. 6a). Additionally, SNPs in GTEx whole-blood eQTL exhibit significantly higher PMP than the background SNPs with matched MAF (Fig. 6b). Similarly, the PMP scores are also highly enriched for SNPs from GWAS catalog compared to other SNPs with matched MAF (Fig. 6c). Together, the results support the utilities of our predictive model in providing a functional prior on common variants to facilitate downstream more focused functional investigation.

**Figure 6:**
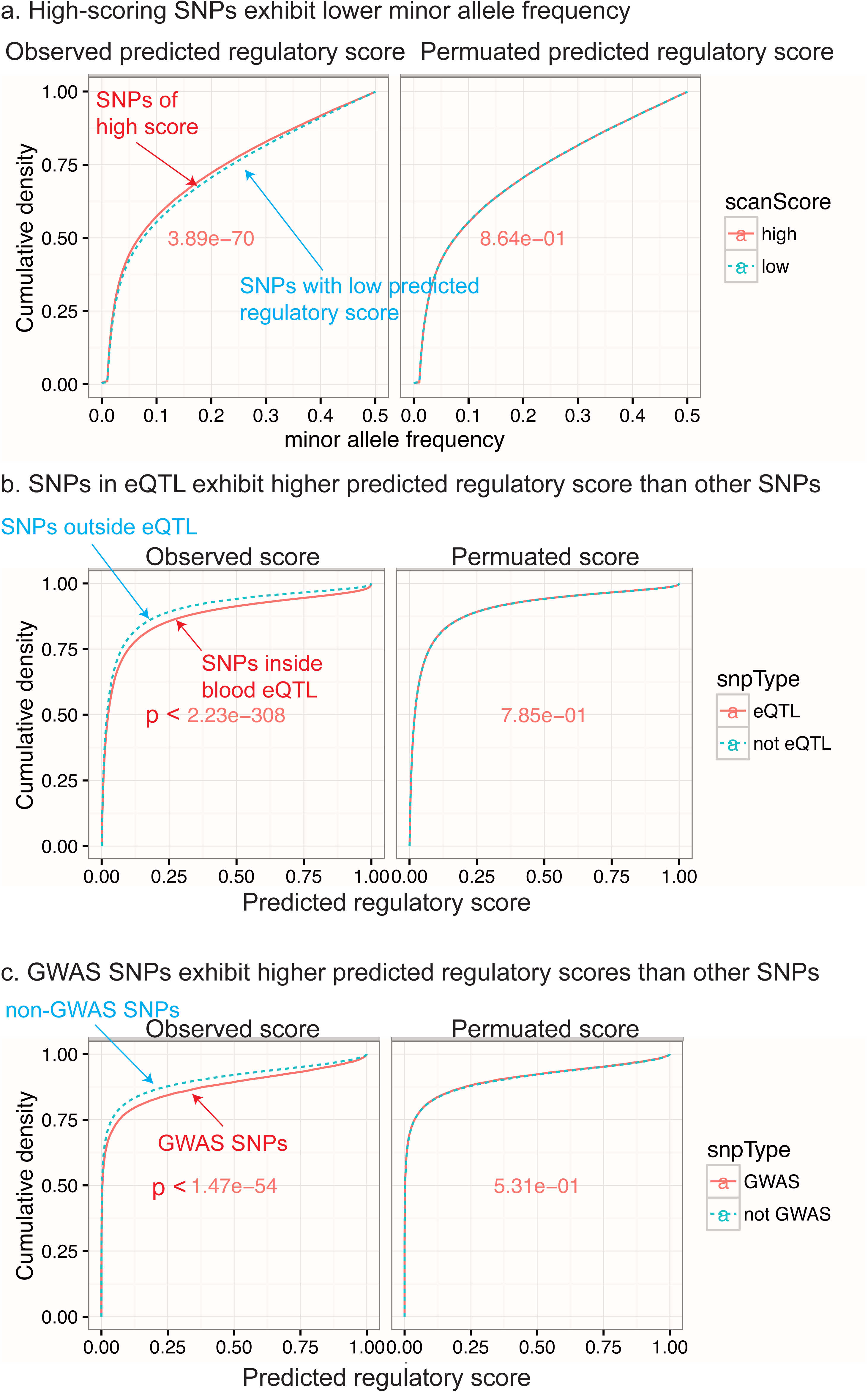
Functional implication of the predicted regulatory scores among the 14.3 million common variants. (a) Distribution of minor allele frequency (MAF). We separated the common variants by variants with high predicted regulatory score (top 1% quantile) and remaining variants. Cumulative density functions of the MAF of the two SNP groups are displayed above. The p-value of the MAF difference between the two SNP groups was calculated using the Wilcoxon rank-sum test, and is shown on each graph. As a control, prediction scores were randomly shuffled among variants and the analysis was repeated, as shown on the right. (b) Functional enrichments for the GTEx eQTL SNPs. We compared the predicted MPRA score of eQTL SNPs in whole-blood with background SNPs with matched MAF. (c) Predicted score of GWAS SNPs. Similar to (b) but comparing SNPs from the GWAS catalog with the remaining SNPs.

## 3 Discussion

MPRA is a promising technology that can efficiently detect thousands of regulatory sequences simultaneously. However, the factors that influence the activity as measured in the assay are not well characterized. In this study, we analyzed two recent MPRA experimental datasets that were generated using naturally occurring eQTLs from human population, in LCL, and based on DNase-sensitive sites and chromatin states in HepG2 and K562 cells. Training machine-learning classifiers to predict regulatory hits from the MPRA data using sequence and functional genomic features, allowed us to uncover meaningful predictive features and achieve promising predictive power. Single-feature tests suggested that one of the most discriminative features for the MPRA data was the presence of DNase-sensitive sites in the regulatory region in question. However, conditioning on DNase-sensitivity in a multiple regression model revealed known sequence motifs, repeat families, and TF binding as the most predictive features based on their coefficients. Most top motifs are active in both HepG2 and K562 cell lines perhaps implying fewer cell-type dependent effects (Fig. 4b). This contrasts with the cell-type specific enrichment of DNase-sensitivity that we observed above.

Although the elasticnet model using functional genomic features performed quite well in most cases, gkm-SVM [28] that uses only the 150-bp sequence information by operating on a gapped k-mer feature space conferred comparable performance. Thus, the regulatory elements that drive MPRA activities can be mostly predicted from the underlying sequence with some small improvements when augmented with functional genomic data. This may be due to the intrinsic properties of regulatory sequence elements as opposed to the dynamic aspects of the regulatory activities implicated in the cell-specific epigenomic data. Another possibility could be that the assays were performed on episomes, which may exhibit different epigenomic landmarks compared to the reference epigenomic annotations obtained from the native chromosomes. Nonetheless, as demonstrated above, we gained biological interpretations by explicitly leveraging the functional genomic features to predict the MPRA outputs than solely based on sequence.

Compared to predicting activating regions, repressive regions were much more difficult to predict, with lower performance among all methods. It is possible that repressive elements may require a specific context or co-factors to have an effect, more so than activating elements. Also, our knowledge database (functional genomic features) for repressive elements is more limited than for enhancer elements.

In terms of testing performance by training on one MPRA dataset and testing on another MPRA dataset, we found that the models using sequence features (gkm-SVM and elasticnet-gkm) compared favorably to the model using only the genomic annotations (elasticnet), implying that the cell-invariant regulatory elements are predominantly implicated in the sequence. Interestingly, models trained on predicting activating (repressive) regulatory hits conferred better or similar prediction accuracy in predicting repressive (activating) regulatory hits in *the same cell type* compared to models trained on predicting activating/repressive regulatory hits in *the other cell type*. This implies that some cell-specific regulatory elements may serve both as activating and repressive sites, which has also been previously observed with a simpler overlapping strategy [22]. The results also imply that different cell types may have different sets of sequence features that dictate regulatory elements. Specifically, the models trained on one cell type performed better in predicting regulatory hits in the same cell types than they did in different cell types. For example, gkmSVM trained on HepG2 rep performed better in predicting HepG2 act than the model trained on K562 rep. In summary, MPRA prediction accuracy is perhaps a reflection of the corresponding MPRA resolution, the amenabilities of the cell types, the MPRA designs, barcoding differences, and the data calibration procedures.

When extending our model to examine the regulatory potential of common variants not tested in the MPRA datasets used, we found that tens of thousands of untested common variants exhibit high regulatory potential, low MAF, and high frequency of being eQTL and GWAS SNPs. Thus, the results imply that (1) there is a certain evolutionary constraint imposed on the putative regulatory variants (indicated by the lower MAF); (2) despite the different nature of the *in-vitro* MPRA experiments, the underlying regulatory elements implicated in the data are consistent with those from the naturally occurring eQTL among the populations; (3) PMP scores can serve as a useful functional prior for risk GWAS variant prioritization. Therefore we suggest specific functional follow-up on these potentially disease-causing but untested common variants, with perhaps a more specific prioritization or re-weighting scheme. On the other hand, we must also cautiously interpret the results due to the small fraction of positive cases (14-23%), which may affect the robustness of the final trained models and the co-linearity existing among the features that we used. With large-scale MPRA data becoming increasing available, similar supervised learning approaches will prove extremely useful in giving insights to experimental outcomes and helping subsequent experimental designs. Thus, we envision our computational approach will serve as a general workflow for future MPRA designs and data explorations.

## 4 Materials and methods

### 4.1 Construction of training data from MPRA experimental data

#### 4.1.1 MPRA data derived from eQTL

The first main training MPRA dataset that we used was testing 3,642 eQTLs discovered by Geuvadis et al., after performing RNA-seq on lymphoblastoid cell lines (LCLs) from individuals of European (EUR) and West African (YRI) ancestry [21]. In total, there were 29,059 150-bp regions that contain either the lead SNPs based on the eQTL analysis or SNPs with perfect linkage-disequilibrium (LD) with the lead SNPs. We formulated the MPRA prediction as a binary classification task. Positive and negative training cases were created as follows. We selected 150-mers with Bonferroni-adjusted p-values < 0.01 based on differential analysis comparing RNA counts with plasmid counts [21]. We then filtered out the 96 sequences with negative fold-change (i.e., RNA counts lower than plasmid counts), resulting in 2,770 “Regulatory hits” hits. The 18,100 150-mers within one standard deviation of the mean fold-change were classified as negative training cases (Fig. 1b).

#### 4.1.2 MPRA tiling data derived from chromatin-states

The other MPRA training datasets were derived from a pool of 15,720 295-bp DNase I hypersensitive sites detected in four cell types (a few of the DNase sites were tiled more than once), namely HepG2, H1hesc, K562, and Huvec, contributing 3,930 sites each [22]. The selection of these sites was based on the 25 chromatin states of the ChromHMM model [22]. Each of the 15,720 295-bp regions was tiled by 31 145-bp oligos with 140-bp overlap (5-bp offset) between consecutive oligos. Slightly fewer than 487,320 sequences (31 × 15, 720 = 487, 320) were successfully synthesized and tested in HepG2 and K562 cells. Sharpr-MPRA was developed to infer the transcriptional activity at a nucleotide resolution, conditioned on the observed MPRA signals, resulting in 4,637,400 data points (15, 720 × 295 = 4,637,400) per cell type [22].

To construct confidence negative (non-regulatory) and positive (activating and repressive) training cases, we fit two 2-component Gaussian mixture models (GMM) on each tiling array dataset to obtain data points belonging to transcriptionally activating and repressive components with high statistical confidence. Specifically, to fit the activating component, we first filtered out repressive regulatory hits by fitting a k-means model with *k* = 3 on the dataset and subsequently removed data points assigned to the cluster exhibiting the lowest mean (i.e., negative mean). We then fit a two-component GMM on the remaining data points with an expectation-maximization (EM) algorithm using the ‘mixtools’ R package [29], where the initial mean and standard deviation of each component were estimated from k-means with the initial mixing proportion equal to 0.5. After the EM algorithm converged within 1e-8, we obtained the lowest value as the activating threshold among the data points belonging to the activating mixture component (i.e., component with the greater Gaussian mean) with posterior probabilities greater than 0.9. Because our dataset was 1-dimensional, selecting data points above this threshold as the positive training examples is equivalent to choosing activating regulatory hits with false discovery rate < 0.1. To choose repressive training examples, we first removed the data points in the above k-means cluster (*k* = 3) with the highest center (i.e., activating cluster) and performed the same procedure as above except that we chose the posterior cutoff to be 0.95 based on the fact that the distribution mode of the tiling data is slightly left shifted from zero (Fig. S1).

To account for the random initialization of the k-means, we repeated the above procedures 500 times and took the average over the thresholds. We then selected the max-scoring nucleotide positions from each 295-bp region based on these thresholds, referred to as “anchors”, with the inferred activities at 90% and 95% posterior confidence corresponding to activating and repressing regions, respectively. We then added 75-bp up-stream and 74-bp down-stream to each anchor, resulting in 1,691 (1,595) and 1,081 (1,872) activating and repressive non-overlapping 150-mers in HepG2 (K562) cells. Finally, the 10,479 (11,401) negative 150-mer negative control examples in HepG2 (K562) were selected from the positions that exhibited activities within 1% standard deviation around but not equal to zero, and not overlapping with the activating or repressive training cases (Fig. 1b). Initial results showed subsequent model performance was not sensitive to the small changes in thresholds (e.g., within 0.5 or 1.5 standard deviation) in defining the negative training cases. Notably, models trained to predict activating and repressive regions used the same negative control training cases.

### 4.2 Single-feature tests

We tested enrichment for the regulatory hits for each feature. Specifically, we performed a hypergeometric test for each binary features using the R built-in function **phyper** and a simple logistic regression for each continuous feature using `glm(y x, family=bionomial, data)`. We calculated p-values from log ratio tests by the `summary` function. Enrichment scores were defined as the - log10 P-value from each test.

### 4.3 Classification models

To construct the MPRA classifiers, we trained a gapped k-mer SVM (gkm-SVM) to predict the regulatory hits using sequence features, and elasticnet using the above-mentioned functional genomic 3,171 features. For each method, we used an existing R package, and default settings were used unless noted otherwise. In particular, we used the R package `gkmSVM` [13, 28] to classify regulatory hits based on all possible 10-mers from the 150-mer training sequences; we used R package `glmnet` [31] with ‘family’ parameter set to “binomial”, alpha to 0.5, and lambda determined by cross-validation (`cv.glmnet`). We also trained a separate “elasticnet gkm” using both the 3,171 functional genomic features and the sequence feature score as fitted by the gkm-SVM model.

### 4.4 Method evaluations

To evaluate each method, we performed N-fold cross-validation. We divided the training datasets into 22 folds, each corresponding to one of the 22 chromosomes. Each method was trained on 21 folds and validated on the remaining fold. Thus, each example was assigned a prediction score when in the validation fold. To evaluate classification performance, we used the R package ROCR [32] to calculate the area under the curve (AUC) of receiver operating characteristic (ROC; AUROC), precision-recall (PRC; AUPRC), and the AUROC up to 5% false positive rate.

### 4.5 Scoring common variants using the MPRA-trained models

We downloaded 14,328,088 common variants from UCSC database (http://hgdownload.soe.ucsc.edu/goldenPath/hg19/database/snp146.txt.gz). We positioned each SNP at the center of a 150-bp (position 76) window to construct the 3,171 features consistent with the featurization procedure described in the main text. We then applied the models trained on each of the full MPRA datasets to score each SNP.

## Supplementary Information

### Supplementary Fig

Figure S1: Tiling MPRA SHARPR score distribution for HepG and K562 cells.

Figure S2: Illustration of fitting 2-component GMM to tiling array data to obtain confidence activating and repressive regulatory training examples.

Figure S3: Predicted regulatory score of common variants in the training MPRA sequences compared with common variants outside of the training MPRA sequences.

Figure S4: Statistical confidence of differential minor allele frequency between high-scoring common variants and low-scoring common variants. We divide the common variants into two groups, one containing SNPs with predicted scores above top 1% quantile and one containing the remaining SNPs. We tested the difference between the two group by Wilcoxon rank-sum test for each of the 15 models (model+training data). Statistical scores displayed in the bar plot are the −log p–values adjusted by Benjamini-Hochberg method. As control, we randomly shuffled the scores from each model and repeated the same analysis (i.e., “_rand”).

Figure S5: We compared the predicted MPRA score of eQTL SNPs in each tissue with background SNPs with matched minor allele frequency. Heatmap color indicates the statistical significance of the −log p–values adjusted for multiple testing by BH-method.

Figure S6: Predicted score of GWAS SNPs. (a) Similar to Fig. 5 but using all of the GWAS SNPs from GWAS catalog. The barplot displays statistical signicance of the −log p–values adjusted for multiple testing by BH-method.

### Supplementary Table

Table S1: All candidate features considered.

Table S2: Cross validation performance.

Table S3: Testing performance.

Table S4: Summary of predicted regulatory variants by each trained model.

Table S5: Predicted MPRA score from 15 models for all of the 14.3 million common variants. Download link http://people.csail.mit.edu/yueli/mpra/TableS5_comVar.gz

